# How does the strength of selection influence genetic correlations ?

**DOI:** 10.1101/2020.06.24.169482

**Authors:** Stéphane Chantepie, Luis-Miguel Chevin

**Affiliations:** Independent Researcher, Antony, France; Centre d’Ecologie Fonctionnelle et Evolutive (CEFE), CNRS, University of Montpellier, University of Paul Valéry Montpellier 3, EPHE, IRD, France

**Keywords:** stabilizing selection, genetic variance-covariance matrix, genetic drift, selection-mutation-drift equilibrium, genetic correlation

## Abstract

Genetic correlations between traits can strongly impact evolutionary responses to selection, and may thus impose constraints on adaptation. Theoretical and empirical work has made it clear that, without strong linkage, genetic correlations at evolutionary equilibrium result from an interplay of correlated pleiotropic effects of mutations, and correlational selection favoring combinations of trait values. However, it is not entirely clear how the strength of stabilizing selection influences this compromise between mutation and selection effects on genetic correlations. Here, we show that the answer to this question crucially depends on the intensity of genetic drift. In large, effectively infinite populations, genetic correlations are unaffected by the strength of selection, regardless of whether the genetic architecture involves common small-effect mutations (Gaussian regime), or rare large-effect mutations (House-of-Cards regime). In contrast in finite populations, the strength of selection does affect genetic correlations, by shifting the balance from drift-dominated to selection-dominated evolutionary dynamics. The transition between these domains depends on mutation parameters to some extent, but with a similar dependence of genetic correlation on the strength of selection. Our results are particularly relevant for understanding how senescence shapes patterns of genetic correlations across ages, and genetic constraints on adaptation during colonization of novel habitats.

## Introduction

Adaptation is inherently a multidimensional problem. Organisms live in complex environments composed of multiple niche axes (Hutchinson, 1957), which exert natural selection on phenotypes composed of multiple traits that get integrated during development (Fisher, 1930). This complexity can limit the process of adaptive evolution. First, the mere fact that multiple traits are under selection can slow down adaptation, which has been described as the cost of complexity (Fisher, 1930; Orr, 2000). And second, genetic correlations between traits can constrain the response to selection for any of these traits, thereby limiting the ensuing increase in fitness by adaptive evolution (Hansen and Houle, 2008; Walsh and Blows, 2009; Lande, 1979; Chevin, 2013; Agrawal and Stinchcombe, 2009; Connallon and Hall, 2018; Etterson and Shaw, 2001). The evolutionary quantitative genetics theory underlying these predictions (Lande, 1979) was soon followed by a related formalism for measuring selection on correlated characters (Lande and Arnold, 1983; Lande, 1979). This has fostered much interest in the last decades for measuring patterns of genetic correlations among traits, in order to quantify constraints on adaptation (reviewed in Agrawal and Stinchcombe, 2009). Such constraints can also be interpreted geometrically (Walsh and Blows, 2009), as genetic correlations can influence the major axis of genetic variation across multiple traits, orienting evolution along lines of least resistance (Schluter, 1996).

Beyond quantifying the consequences of genetic correlations on rates on adaptation, understanding what shapes constraints on adaptation ultimately requires investigating the factors that govern the evolution of the **G** matrix, which includes all the additive genetic variances of traits and covariances among traits (Lande, 1979). This has been a topic of intense research, both theoretically and empirically. Theoretical work has made it clear that genetic correlations evolve in response to (i) correlated pleiotropic mutation effects on traits, and (ii) correlational selection favoring combinations of trait values between pairs of traits (Lande, 1980; Turelli, 1985). Random genetic drift may also play an important role (Jones et al., 2003), but this was mostly investigated through individual-based simulations, and few analytical results exist to guide intuition in that respect. In addition, patterns of environmental change (Jones et al., 2004, 2012) and epistatic interactions among loci (Jones et al., 2014) can also influence the shape of the **G** matrix and evolution of genetic correlation, but we will not address them here. On the empirical side, it was recently demonstrated that the genetic divergence of multiple traits across several *Drosophila* species is aligned with the major axis of both the **G** matrix of additive genetic variation within species, and the **M** matrix of mutation effects on these traits (Houle et al., 2017). Natural selection was not measured in that study, but another study on the same set of traits has demonstrated that their genetic correlations can evolve in response to experimental patterns of correlational selection (Bolstad et al., 2015).

Since genetic correlations result from a compromise between mutational correlations and correlational selection, we may wonder: how do they change as the strength of selection varies? And more generally: how does the overall shape of the **G** matrix change as a fitness peak becomes broader (thus causing weaker selection), or narrower (stronger selection), while keeping the same overall shape (as illustrated in Fig. 1a)? This simple question has received surprisingly little attention, despite its general importance in evolutionary biology. In particular, it bears on our understanding of the evolution of senescence by mutation accumulation, whereby relaxed selection on later age classes allow for accumulation of more genetic variance of traits (Charlesworth and Hughes, 1996). A multivariate extension of this argument might suggest that the **G** matrix becomes more similar to the mutation **M** matrix in older ages, because they undergo relaxed selection. However, the premises that underlie this argument have yet to be explored more thoroughly.

**Figure 1:**
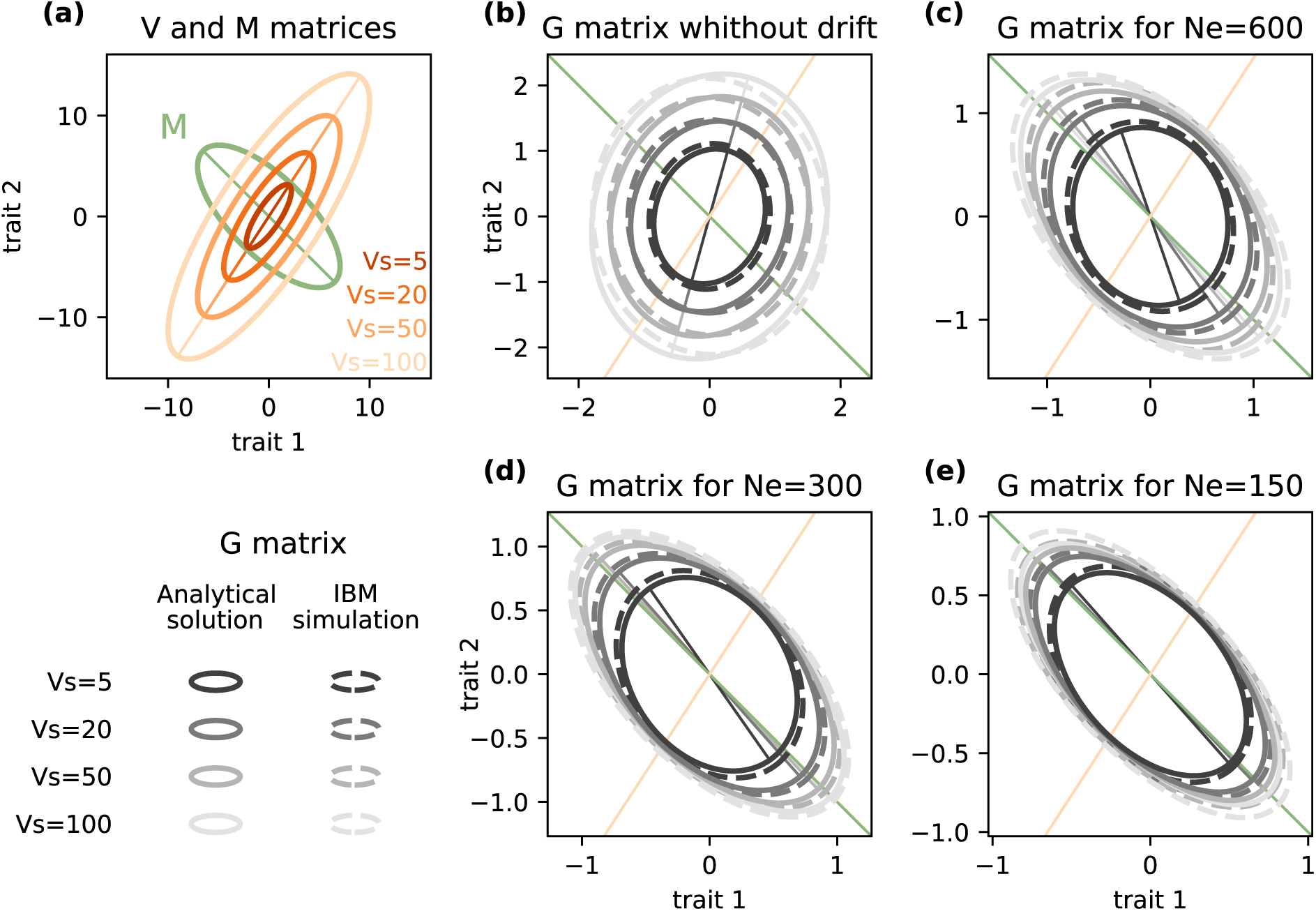
Influence of selection strength and genetic drift on the G matrix (Gaussian regime). (a) Orientation and shape of the mutation matrix **M**, and selection matrix **V** with variable selection strengths. First eigenvectors are represented with colored lines. (b-e) Shape and orientation of the **G** matrix as the width of the fitness peak *V*_*s*_ varies. Solid ellipses (along with their first eigenvectors) represent the analytical predictions from equation (7) that neglects genetic drift in (b), or equation (8) that accounts for genetic drift in (c-e). Dashed ellipses show the mean estimates from IBM simulations with *N*_*e*_ = 5000 (b), 600 (c), 300 (d) and 150 (e). Parameters used: *n* = 50, *µ* = 0.01, *ρ*_*m*_ = *−*0.7, *ϕ*_*m*_ = 1, *V*_*α*_ = 0.0025 and *ρ*_*s*_ = 0.8, *ϕ*_*s*_ = 2, and *V*_*s*_ = 5, 20, 50, 100.

Here, we investigate theoretically how the overall strength of selection influences evolution of genetic correlations, and the shape and orientation of the **G** matrix. Using analytical results and individual based simulations, we show that the relative importance of mutation vs selection in shaping the **G** matrix critically depends on random genetic drift.

## Methods

### Model

As in standard quantitative genetic models, we assume that the multivariate phenotype **z** can be partitioned into a breeding value determined by the genotype, plus a residual component of variation (often described as the environmental component), which is normally distributed with mean 0 and covariance matrix **E**. In each generation, mutations occur with probability *µ* at each allele of *n* diploid loci, such that the total mutation rate is 2*nµ*. Mutation increments the phenotypic value at the mutated allele by an effect that is unbiased (does not change the average breeding value), but can change the genetic (co)variances between traits. Specifically, we assume multivariate normally distributed mutation effects ***α***, with mean 0 and the same covariance matrix **M** at each locus, which we parameterize (for two traits) as

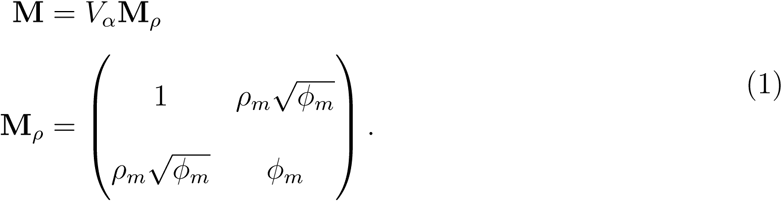

The parameter *V*_*α*_ is the variance of mutation effects on trait 1 (a scalar), *ϕ*_*m*_ is the ratio of mutational variances between traits 2 and 1, and *ρ*_*m*_ is the mutational correlation. When *ϕ*_*m*_ = 1 the two traits have the same mutational variance, and **M**_*ρ*_ is a mutational correlation matrix. The multivariate phenotype is under stabilizing selection towards an optimum phenotype ***θ***, which we assume constant for simplicity. This is modeled as classically by letting the fitness of individuals with multivariate phenotype **z** (relative to the fitness of the optimum phenotype) be

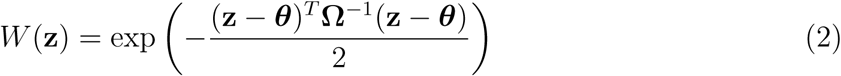

where the matrix **Ω** determines the breadth and orientation of the fitness peak. Averaging over the distribution of residual phenotypic variation, the fitness function on breeding values **x** (relative to the fitness of the optimum breeding value), which determines evolution of the **G** matrix, is

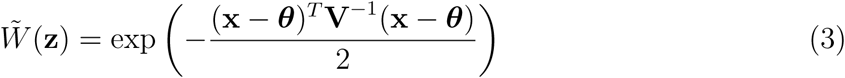

where **V = Ω + E** is the stabilizing selection matrix. Note that because of residual non-heritable phenotypic variation, the absolute fitness of individuals with a given breeding value **x** is actually reduced, by a factor 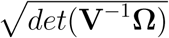 (where *det* denotes the determinant of a matrix), relative to that of individuals with the same realized phenotype **z** = **x**. This amounts to reducing the effective number of parents in the population: even in a Wright-Fisher population of size *N*, the effective size that matters for random genetic drift (change in the distribution of breeding values) is in fact 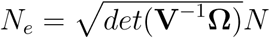 (and in a non-Wright-Fisher population, *N* should be replaced by the effective size operating at the level of the expressed phenotypic trait). In other words, selection on non-heritable phenotypic variation increases the intensity of genetic drift on heritable phenotypic variation. This fact, which was largely overlooked in previous studies on this topic (e.g. Lande, 1976, 1979; Burger et al., 1989), becomes important under strong selection and low population size.

The selection matrix **V** can be written similarly to **M** as

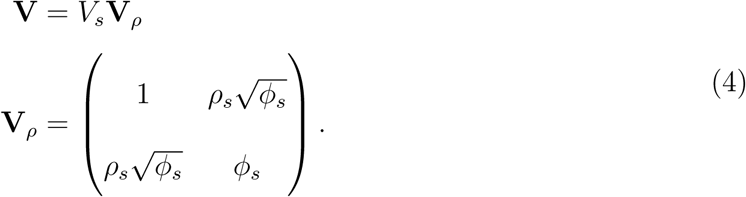

The scalar *V*_*s*_ determines the width of the fitness peak on breeding values, and is inversely proportional to the strength of stabilizing selection, while *ϕ*_*s*_ controls the ratio of strengths of selection between the two traits. The selective correlation *ρ*_*s*_ determines what genetic correlation is favored by natural selection. Figure 1a illustrates how these parameters translate into the shapes of the mutation and selection matrix.

### Individual-based simulations

We tested the accuracy of expected **G** matrix and genetic correlation at mutation-selection(-drift) equilibrium using individual-based, genetically explicit simulations. The traits were determined by *n* = 50 unlinked diploid loci, with alleles assumed to be fully pleitropic (i.e. affecting all the phenotypic traits under selection). We simulated populations of hermaphroditic, sexually reproducing individuals with non-overlapping generations. The life cycle included three steps:

1. **Computing the phenotype and fitness for each individual**. For each trait, the phenotype was estimated by summing breeding values across all loci and alleles. A residual component of variation was then added for each trait, with mean 0, variance 1, and no covariance between traits. The expected fitness of each individual was then computed based on their phenotypic values and the stabilizing selection matrix **Ω**, as defined by equation (2). The fitness optimum was arbitrarily set to zero.
2. **Sequential reproduction based on reproductive fitness**. Two adults were drawn randomly with a probability equal to their expected fitness and mated to produce exactly one offspring. Selfing was not allowed, and being involved in a reproductive event did not change the probability to mate again. The sequence was repeated *N* times.
3. **Offspring production**. The genotypes of offspring were produced by drawing one random allele from each parent at each locus, thus modelling fully unlinked loci. The probability that a mutation occurred a each allele of each locus was *µ*. Mutation had additive effect on the traits, modifying the phenotypic value of the mutated allele by an amount drawn from a multivariate normal distribution, with means zero (unbiased mutation) and mutation variance-covariance matrix **M**.

This life cycle, developed by Revell (2007), ensures that the population size *N* is constant and equal to the effective population size *N*_*e*_ in the absence of selection. Here, to account for the reduction in effective population size caused by selection on the residual component of variation (see previous section), for each required value of *N*_*e*_ we used 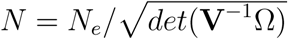 as the population size in the simulations. Since all our formulas depended on the matrix **V** of selection on breeding values (eq. (3)), rather than the matrix **Ω** for selection on the expressed phenotype (eq. (2)), we parameterized simulations in terms of **V**, and then transformed them to **Ω** before starting the simulation using **Ω** = **V** *−* **E** (as per eq. (3)), where **E** = **I** under our assumption of uncorrelated environmental effects with variance 1.

Individual-based simulations were all run over 50000 generations. To ensure that the expected genetic covariance matrix 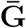 was estimated after a pseudo-equilibrium is reached (i.e., at stationarity), only the 30000 last iterations from the chain were used to estimate the mean.

## Results

### Without drift, genetic correlations are unchanged by selection strength

Using similar assumptions as here, Zhang and Hill (2003) showed that in an infinite population, and in the limit of rare mutations of large effect (so-called House-of-Cards regime, HoC below; Turelli, 1984, 1985; Bulmer, 1989; Bürger, 2000; Johnson and Barton, 2005), the genetic correlation *ρ*_*G*_ between two traits at mutation-selection balance with weakly linked loci is

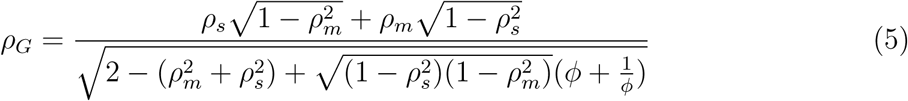

where 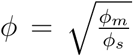. Remarkably, this shows that genetic correlations at mutation-selection balance depend neither on the absolute strength of stabilizing selection 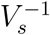 nor on the magnitude of mutational variance *V*_*α*_, but instead on the ratio of strengths of stabilizing selection between the two traits, times the ratio of their mutation variances (summarized by the compound parameter *ϕ*). This means that narrowing the fitness peak, thereby increasing the strength of stabilizing selection on all traits, does not tilt the balance of genetic correlations *ρ*_*G*_ towards selective correlations *ρ*_*s*_ and away from mutational correlations *ρ*_*m*_, as long as the overall shape of the mutation and selection matrices do not change. The same is true of increasing the mutational variance and covariances of all traits by the same factor. In the special case where *ϕ* = 1, such that the the ratio of strengths of stabilizing selection on the two traits equals the ratio of their mutational variances, equation (5) further simplifies as

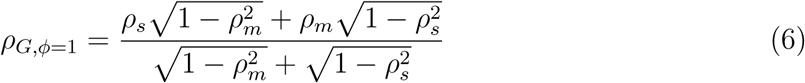

which only depends on mutational and selective correlations.

This result was obtained under the HoC regime, which is known to have different properties from a regime of common mutations of weak effect, known as the Gaussian regime (Kimura, 1965; Lande, 1976; Bulmer, 1989; Bürger, 2000; Johnson and Barton, 2005). But in fact, genetic correlations also do not depend on the strength of selection under the Gaussian regime. To see this, we rewrite the equilibrium for the **G** matrix derived by Lande (1980) under the Gaussian regime, replacing **V** and **M** with their expressions in equations (1) and (4), to get 

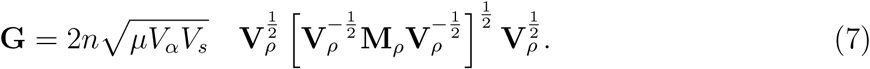

The first scalar term is the same as for the genetic variance of a single trait at mutation-selection balance in this regime (Kimura, 1965; Lande, 1976). The second term is a matrix that captures all the features of **G** matrix shape, including genetic correlations. Equation (7) shows that changing the overall strength of selection 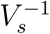, or the scale of mutational variance *V*_*m*_, only magnifies or shrinks the **G** matrix, but does not change genetic correlations in any way, nor any other aspect of **G** matrix shape.

Figure 1 shows examples of **G** matrices under variable strength of stabilizing selection (where the fitness peak becomes broader, and selection becomes weaker, as *V*_*s*_ increases, Fig. 1a), in the Gaussian regime. Continuous ellipses in Figure 1b represent the analytical prediction for **G** from equation (7), while dashed ellipses show results from genetically explicit individual-based simulations (IBM) using the same parameters, but finite population size *N*_*e*_ = 5000. The prediction that the strength of stabilizing selection does not affect the orientation of the **G** matrix in a infinite population is already close to holding in simulations with *N*_*e*_ = 5000. The volume of the **G** matrix increases, but its orientation and shape change little as the strength of selection decreases. The resulting genetic correlation is also little influenced by the strength of selection in simulations with *N*_*e*_ = 5000 (dark blue dots in Figure 2b), and remains close to the expected compromise between the mutational and selective correlations predicted by equation (5) (black line in Figure 2b), which does not depend on *V*_*s*_.

**Figure 2:**
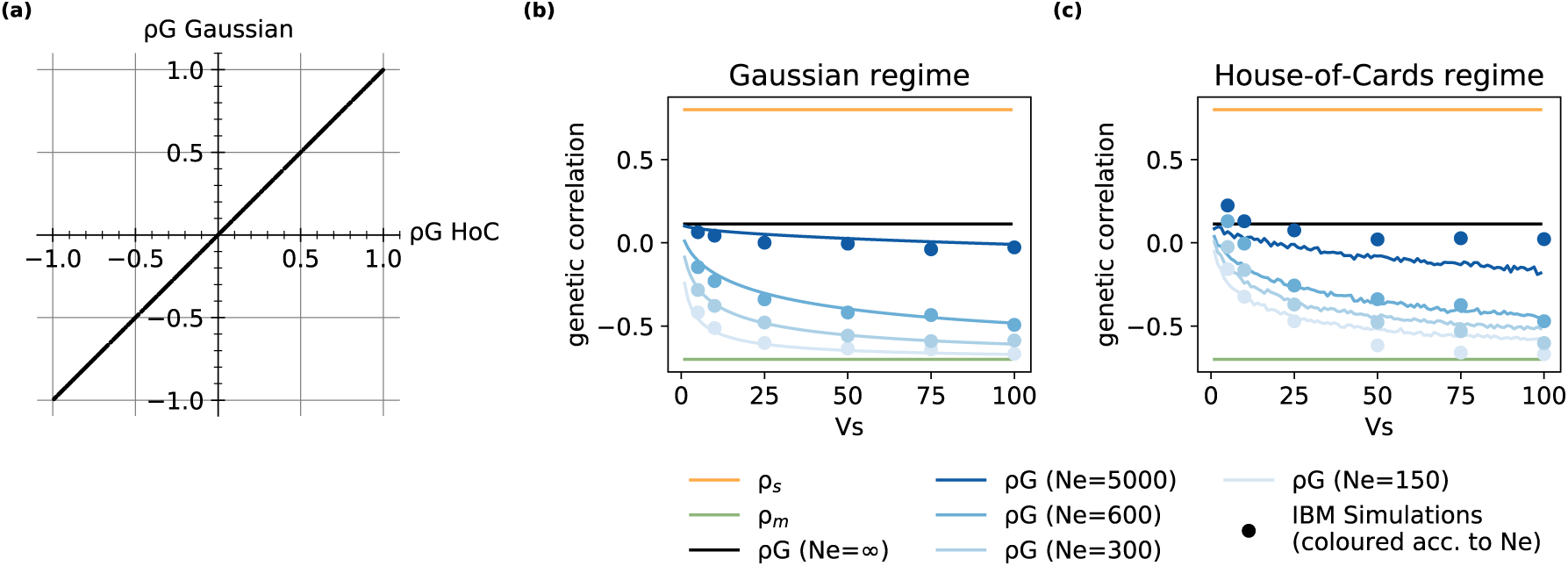
Influence of selection strength and genetic drift on genetic correlations under different mutation regimes. (a) The genetic correlation in an infinite population, at equilibrium between selection and abundant mutations of weak effects (Gaussian approximation, from eq. (7)) is plotted against its expectation under rare mutations of large effects (house-of-cards approximation, eq. 5), for 500 random pairs of mutation **M** and selection **V** matrices. (b-c) The genetic correlation is plotted against the width of the fitness peak *V*_*s*_, for different effective sizes *N*_*e*_. The parameters values in (b) are the same as in Figure 1, corresponding to the Gaussian mutation regime. In (c), the mutation parameters are instead *V*_*α*_ = 0.05 and *µ* = 0.0002, corresponding to the House-of-cards regime. Blue points correspond to the genetic correlation simulated with individual-based models (IBM). The black line represents the analytical expectation without drift (eq. (5)) in both cases. The blue lines represent the analytical prediction with drift (eq. (8)) in (b), and expectations over 10000 randomly drawn mutation effects *α* (from eq. (9)) in (c).

Even though the Gaussian and Hoc regime have very different properties in terms of the maintenance of genetic variance for each trait (Turelli, 1985; Bürger, 2000), they strikingly lead to the same genetic correlation among traits in an infinite population. This was already suggested in numerical explorations by Turelli (1985), but we confirmed this here more extensively. In particular, when *ϕ*_*m*_ = *ϕ*_*s*_ = 1, such that the mutation and selection matrices are both proportional to correlation matrices (with only 1 on the diagonal), then deriving the genetic correlation in the Gaussian case from the **G** matrix in equation (7) leads to equation (6). In the more general case, an analytical formula also exists for the genetic correlation based on equation (7), but it is unwieldy. Instead of comparing Gaussian and Hoc formulas for genetic correlation, we drew random matrices **V** and **M** from a Wishart distribution (with expectation **I**), a natural distribution for covariance matrices, which allows variance and covariance terms to vary randomly. We then computed the expected **G** matrix under the Gaussian regime (from eq. (7)), from which we extracted genetic correlations, which we then compared to equation (5). Figure 2a shows that the genetic correlation under the Gaussian regime, which assumes frequent mutations of small effects, is perfectly predicted by that under the House-of-Card regime, which instead assumes rare mutations of large effects. The blue dots in Figure 2c show genetic correlations for different values of the selection parameter *V*_*s*_, estimated from individual-based simulations with parameters that correspond to the HoC regime. These correlations are very similar to those in the Gaussian regime in Figure 2b, and close to their expectation in eq. (5).

In short, genetic correlations at mutation-selection balance do not change with the overall strength of stabilizing selection, and this conclusion holds generally across a broad range of mutation and selection parameters, spanning different evolutionary regimes. So should we then conclude that correlational selection always has the same influence on genetic correlations, and never becomes dominated by the influence of mutational correlations, even as the strength of selection becomes vanishingly small?

### Drift controls the balance between mutation and selection’s effects on genetic correlations

In fact, genetic correlations may indeed become more similar to mutational correlations as the strength of selection decreases, but only in the presence of random genetic drift. In a population with finite effective size *N*_*e*_, random genetic drift causes a reduction in heterozygosity, and thus in additive genetic variance, by a proportion 2*N*_*e*_ per generation. Accounting for this effect, we found that the expected **G** matrix at mutation-selection-drift equilibrium in the Gaussian regime is (Appendix A1)

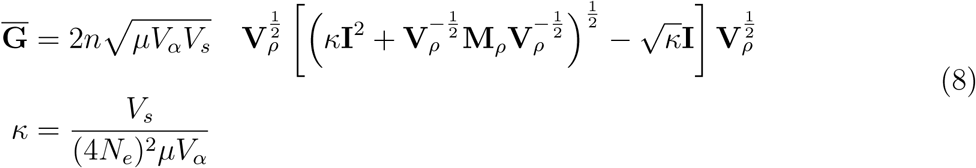

where 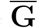 denotes an expectation over the stochastic evolutionary process (because of random genetic drift). As in equation (7), the first scalar term in equation (8) is the same as for the genetic variance of a single trait at mutation-selection balance in this regime (Kimura, 1965; Lande, 1976), while the matrix product determines **G** matrix shape. Equation (8) shows that a single compound scalar parameter, *κ* = *V*_*s/*_[(4*N*_*e*_)^2^*µV*_*α*_], determines how the orientation and shape of the expected **G** matrix change under mutation, selection, and drift (since elements of **V**_*ρ*_ and **M**_*ρ*_ scale on the order 1 by construction). When *V*_*s*_ *≪* (4*N*_*e*_)^2^*µV*_*α*_ (*κ* very small), the mutation rate and mean selection coefficient of new mutations are both large relative to the intensity of drift (proportional to 1*/N*_*e*_), so genetic correlations are mostly determined by mutation and selection, with little influence of genetic drift. In the limit *κ →* 0, equation (8) tends to the mutation-selection balance in equation (7). In contrast, drift dominates when *V*_*s*_ ≫ (4*N*_*e*_)^2^*µV*_*α*_ (*κ* very large), and the expected **G** matrix then becomes increasingly similar to the mutation matrix **M**. This can be seen when comparing **G** matrices in panels b to e in Figure 1, as well as genetic correlations for different darknesses of blue in Figure 2b. Furthermore for a given *N*_*e*_, the genetic correlation and orientation of the **G** matrix become more similar to those of mutation as the strength of selection decreases (increasing *V*_*s*_, lighter ellipses in Fig. 1b-e, and rightmost values in Fig. 2b).

We have shown above that the type of mutation-selection regime (HoC vs Gaussian) does not influence genetic correlations in an infinite population (Fig. 2a), but is it also the case in a finite population with substantial genetic drift? For the HoC regime, the expected **G** in an infinite population is proportional to the expectation of 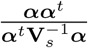 over the distribution of mutation effects ***α*** (Zhang and Hill, 2003, eq. 2). In a finite population, accounting for the reduction in heterozygosity caused by both stabilizing selection and random genetic drift, this approximation becomes (adapted from Burger et al., 1989, “stochastic house of cards” regime)

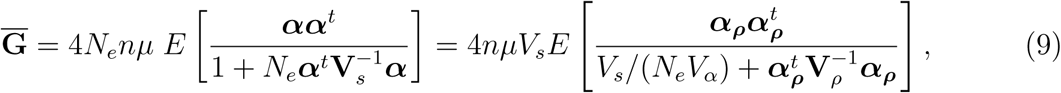

where *E*[] denotes an expectation over the distribution of mutation effects, and 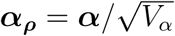 are scaled mutation effects, with covariance matrix **M**_*ρ*_ as per equation (1). Analogously to equations (7) and (8), the first scalar term in equation (9) equals the equilibrium genetic variance for a single trait in the HoC regime, while the expectation includes all the parameters that determine **G** matrix shape. Since elements of **V**_*ρ*_ and **M**_*ρ*_ scale on the order 1 (and hence so do 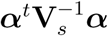 and elements of ***αα***^*t*^), the scalar parameter *V*_*s*_/ (*N*_*e*_*V*_*α*_) alone determines whether the **G** matrix is more influenced by mutation, selection, or drift. When *V*_*s*_ *≪ N*_*e*_*V*_*α*_, drift can be neglected and the **G** matrix has the same shape as in the HoC equilibrium; in particular, the genetic correlation between two traits is given by equation (5), and is thus the same as in the Gaussian regime. In contrast when *V*_*s*_ *" N*_*e*_*V*_*α*_, drift dominate and the **G** matrix is proportional to **M**, with correlation *ρ*_*m*_.

The stochastic house of cards approximation to the genetic correlation (eq. 9) is somewhat less accurate at predicting results from individual-based simulations than our stochastic Gaussian approximation in the corresponding regime (eq. 9, Fig. 2c, Fig. S2). However, it does capture a similar pattern, where the genetic correlation tends more rapidly towards the mutational correlational with decreasing strength of selection when the effective population size is smaller.

In summary, accounting for genetic drift in finite populations, the genetic correlation spans the same range in all mutation regimes, ranging from the mutational correlation *ρ*_*m*_ when drift dominates, to the compromise between mutation and selection in equation (5) when selection dominates. The mutation regime (Gaussian vs HoC) only determines how the realized genetic correlation interpolates between these two limit cases. In particular, the selection strength 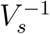 at which genetic correlations transition from being drift-dominated to selection-dominated is multiplied by (16*N*_*e*_*µ*)^*−*1^ in the Gaussian regime relative to the HoC regime (eqs.(7) and (9)). When 16*N*_*e*_*µ <* 1, this means that stronger selection is required to overcome the influence of drift when mutations are abundant but with small effects (Gaussian regime) as compared to rare but with larger effects (and vice versa when 16*N*_*e*_*µ >* 1). But beyond these changes in the quantitative dependence on the strength of selection (which relate to previous findings for a single trait, Hermisson and Wagner, 2004; Bürger, 2000), the qualitative relationship between genetic correlations and the strength of selection given the effective population size remains the same across mutation regimes (Fig. 2b-c).

### Shape, orientation, and correlation

Changes in the relative importance of selection versus genetic drift can influence the shape of the **G** matrix (determined by its eigenvalues), its orientation (determined by its eigenvectors), or both. Any of these effects can translate into changes in genetic correlations, since the latter are only a summary of **G**, which depend on both variances and covariances (and on both eigenvectors and eigenvalues). In particular, genetic correlations may change despite little change in orientation, and no rotation of the axes of the **G** matrix. This is illustrated in Fig. 3, which focuses on the special case where **V** and **M** have the same eigenvectors. When this holds, the eigenvectors of **G** are identical to those of **V** and **M**, as demonstrated in the Appendix A2. This means that the **G** matrix does not rotate when changing the relative importance of drift versus selection; all that changes are the amounts of variation (eigenvalues) along the different axes (eigenvectors). Nevertheless, the genetic correlation still changes according to eq. (9) in this example. For instance, when one eigenvalues becomes dominant, the **G** matrix becomes increasingly elongated along one of the eigenvectors (Fig. 3a, Fig. S1), which translates into larger values of genetic correlations (Fig. 3b). In the more general case where **V** and **M** have different eigenvectors, then changing the relative importance of selection vs drift causes both elongation and rotation of **G** (changes in shape and orientation), but without necessarily causing larger changes in genetic correlations.

**Figure 3:**
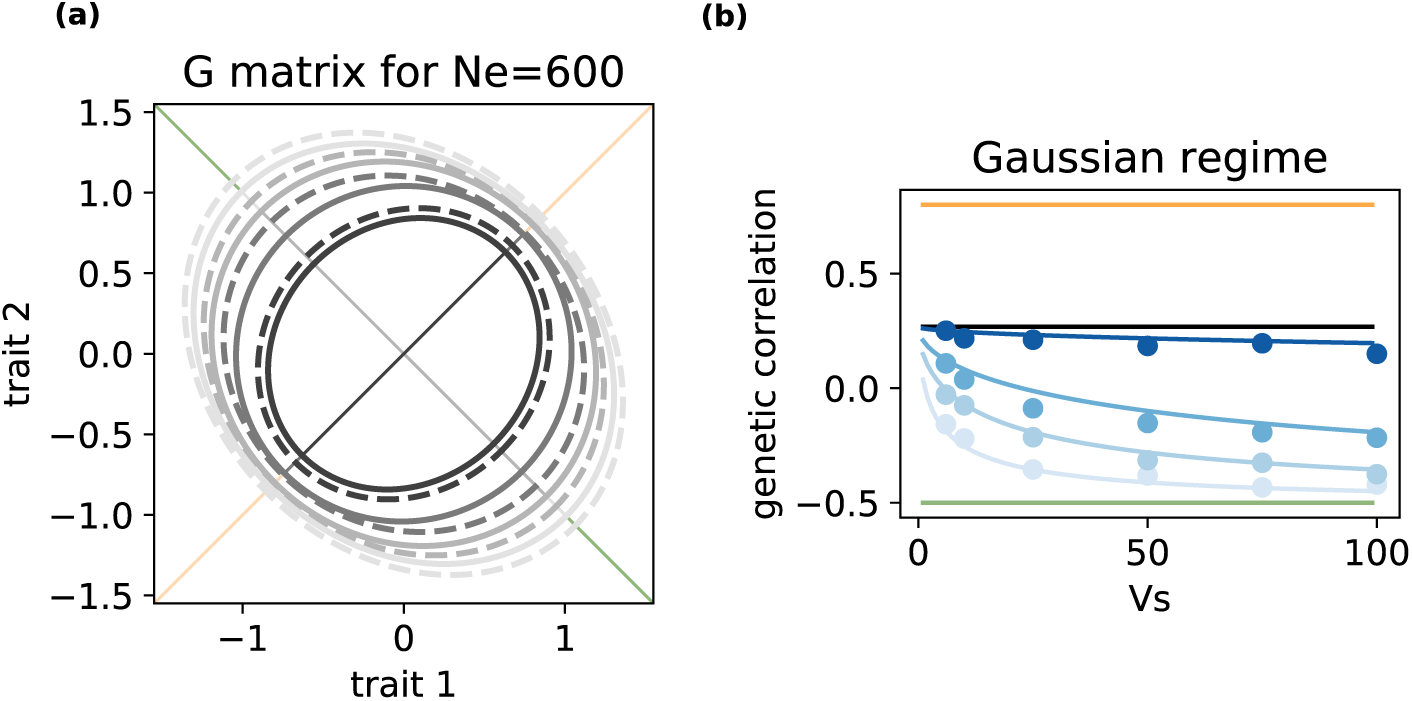
Influence of selection strength and genetic drift on genetic correlations in the special case where V and M have the same eigenvectors. (a) Shape and orientation of the **G** matrix as the width of the fitness peak *V*_*s*_ varies. In this illustrative example where *N*_*e*_ = 600, the first eigenvector of the **G** matrix (plain lines, from gray to black) is orientated either fully along the major axis for selection (eigenvector of **V**, orange line) or for mutation (eigenvector of **M**, green line), depending on the selection parameter *V*_*s*_. (b) The genetic correlation is plotted against the width of the fitness peak *V*_*s*_, for different effective sizes *N*_*e*_ as in Figure 2. Note that, although axes of the **G** matrix do not rotate, the influence of selection strength and genetic drift on genetic correlations is comparable to the general case in Figure 2. Parameters are identical to Figure 1, except that *ϕ*_*s*_ = *ϕ*_*m*_ = 1 such that **V** and **M** have the same eigenvectors, *ρ*_*m*_ = *−*0.5, and the strongest selection is for *V*_*s*_ = 6 instead of 5.

## Discussion and Conclusion

To what extent are genetic correlations between traits shaped by natural selection, or imposed by mutation? This question, which in essence traces back to the debate between mutationists and selectionists in the early days of genetics (recently revived in the light of molecular evidence, Nei, 2013), has received considerable attention from evolutionary biologists. In particular, evolutionary quantitative genetic theory has made it clear that, when phenotypic (co)variances arise from an equilibrium between mutation and stabilizing selection, then genetic correlations are a compromise between the correlation of pleitropic mutation effects on traits, and correlational selection favoring combinations of traits (Lande, 1980; Lande and Arnold, 1983; Jones et al., 2003; Turelli, 1985). The latter can be related to the orientation and elongation of the fitness landscape relating the traits to fitness (as illustrated in Fig. 1a). However beyond this shape of the fitness landscape, how does the overall strength of selection (size of the ellipses in Fig. 1a) influence genetic correlations between traits? As selection becomes weaker, the fitness peak becomes flatter, with a broader range of phenotypes having equivalent fitness, so does that reduce the influence of selection on genetic correlations? Perhaps surprisingly, the answer is no in an effectively infinite population, which in our simulations was already close to holding for a moderate population size of *N*_*e*_ = 5000 individuals. Strikingly, the same compromise between mutation and selection effects on genetic correlations holds regardless of the strength of selection, and regardless of whether genetic (co)variances are caused by common mutations of small effect (Gaussian regime, Kimura, 1965; Lande, 1980), or rare mutations of large effect (House-of-cards regime, Turelli, 1984, 1985).

However, the strength of selection starts to matter as the effective populations size *N*_*e*_ becomes smaller, as random genetic drift plays a larger role. The reasons is that, for a given *N*_*e*_ the strength of selection shifts the balance between drift-dominated and selection-dominated evolutionary dynamics. Since genetic correlations equal mutational correlations *ρ*_*m*_ in the former domain, but a compromise between mutation and selection (eq. (5)) in the latter, overall mutation has a stronger influence on genetic correlations than selection (Fig. 2). Previous analyses of **G** matrix evolution under mutation and correlational selection in finite populations has mostly focused on the effect of drift on the stability of the **G** matrix over evolutionary time (Jones et al., 2003), and largely overlooked the influence of drift on the expected **G**. In fact, this influence can be substantial, as shown here; in particular, it determines how the strength of selection affects genetic correlations.

Genetic correlations are often described as a constraint on adaptation (Etterson and Shaw, 2001; Agrawal and Stinchcombe, 2009; Connallon and Hall, 2018), but this need not be true, depending on how the orientation of the **G** matrix relates to that of directional or fluctuating selection in a changing environment (Gomulkiewicz and Houle, 2009; Chevin, 2013; Duputié et al., 2012). In a constant environment as assumed here, the extent to which genetic correlations constrain adaptation depends on how the **G** matrix aligns with the matrix of correlational selection, represented in Figure 1a. Our analytical and simulations results show that genetic correlations, and the overall **G** matrix shape, differ more from those favored by correlational selection at lower effective population sizes. Since **G** becomes more similar to the mutation matrix **M** in this case, this could be interpreted as a mutational constraint on evolution (Nei, 2013). However, this alignment with mutation effects occurs because of a prevalence of genetic drift, which is in fact the main constraint on adaptation in this case, also causing temporal fluctuations in the mean phenotype (Lande, 1979) and the G matrix itself (Jones et al., 2003), and apparent fluctuating selection (Chevin and Haller, 2014).

Our analytical results for genetic correlations and the **G** matrix at mutation-selection-drift balance in the Gaussian regime (eq. (8)) are valid under frequent mutation (Kimura, 1965; Lande, 1980; Bürger, 2000), and we accordingly used high mutation rates in the corresponding simulations. However, note that the nature of loci is not explicit in this model, but in any case these do not represent single nucleotides or even genes. Rather, they represent large stretches of effectively non-recombining portions of the genome, which may influence the traits by mutation. Since free recombination is also assumed across these loci (consistent with most previous studies), the latter can even be thought of as small chromosomes, for which mutation rates of the order to 10^*−*2^ seem reasonable. In addition, we also present theoretical and simulation results at much lower mutation rates (House-of-Cards regime), which lead to similar findings. We assumed universal pleiotropy, whereby all loci have the same distribution of mutation effects on all traits. An interesting extension may be to allow for modular mutation effects, or restricted pleiotropy, whereby each locus can only modify a subset of traits by mutations (Chebib and Guillaume, 2017; Chevin et al., 2010), to investigate whether the mutation regime has a stronger effect on mutation correlations in these scenarios. In terms of selection, we considered a fitness peak with an optimum, in line with most theory on the topic, but genetic correlations can also be favored by other forms of selection, notably disruptive selection (Bolstad et al., 2015), or negative frequency dependence caused by individual interactions (Mullon and Lehmann, 2019), which may lead to different dependencies of genetic correlations on the strength of selection and genetic drift.

Our results lead to some predictions about how genetic correlations should change along lifetime. For traits with ontogenic trajectories, or traits involved in senescence, the phenotypic value at different ages can be considered as different character states, partly controlled by different loci. This underlies both the mutation accumulation theory of senescence applied to quantitative traits (Charlesworth and Hughes, 1996), and the theory of evolution of growth trajectories (Kirkpatrick and Lofsvold, 1992). If the shape of correlational selection does not change with age, then we only expect a reduced strength of selection in older age, because these ages have a smaller reproductive value and hence contribute less to fitness (Lande, 1982; Charlesworth, 1993). We would then predict that genetic correlations should lean more towards mutational correlations in older ages, but only when the effective population size is small, while genetic correlations should remain largely unchanged along lifetime in large populations. This pattern can be investigated by measuring genetic correlations among primary traits (not direct components of fitness) across ages, for different species that differ in effective population size (as estimated by e.g. their molecular polymorphism level).

More broadly speaking, we expect mutational correlations to impose more constraints on evolutionary trajectories in situations where the population size has been reduced, such as bottlenecks during colonization of novel habitats. Since these situations are also likely to be associated with strong directional selection, this should represent a double challenge for colonizing species. Nevertheless, the extent to which mutational correlations *per se* impede responses to directional selection is unclear. Even when genetic correlations are largely shaped by correlational selection (rather than just by mutation), they may still constrain adaptation, if directional selection in a novel or changing environment does not align with the shape of the fitness peak (Chevin, 2013). In any case, our clear delineation of when, and how much, the strength of selection influences genetic correlations, should provide guidelines for analyzing and interpreting genetic constraints on adaptation in the wild.

## Acknowledgments

SC wish to thank Thomas Hansen, David Houle and Christophe Pélabon for discussions on preliminary results. The authors declare no conflicts of interest. This work was supported by the European Research Council (Grant 678140-FluctEvol).

## Author Contributions

SC and LMC conceived the study. SC developed analytical solutions for the Gaussian regime. LMC developed analytical solutions for the House-of-Cards regime. SC developed and performed individual-based simulations. SC and LMC wrote the manuscript.

## Data Accessibility

The C++ software developed and used to perform genetically explicit individual-based simulations will be available under GPL licence on Github after acceptance.

## A Appendix: How does the strength of selection influence genetic correlations ?

### A.1 Steps for solving the mutation-selection-drift equilibrium

By building on the Gaussian approximation of the continuum-of-allele model (Kimura, 1965) and assuming infinite-sized population with non-overlapping generations, Lande (1980) derived a simple expression for genetic covariance matrix equilibrium (**G**) selection-mutation balance. We here extend this result to allow for random genetic drift. At equilibrium, the production of genetic variance due to new polygenic mutations is balanced by the loss of genetic variance to due both stabilizing selection and random drift (Lande, 1979). For a single haploid locus we obtain:

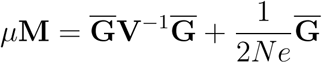

where 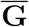 denotes an expectation over the stochastic evolutionary process (because of random genetic drift).

Introducing 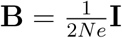 and **U** = *µ***M**, this becomes

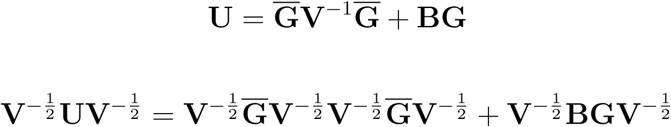

By defining 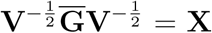 and 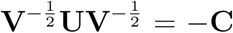, we finally have to solve the quadratic matrix equation:

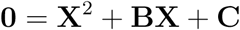

A quadratic matrix equation can be solved explicitly if the following requirements are met: (i) **X**^2^ is preceded by an identity matrix, (ii) **B** commutes with **C**, and (iii) **B**^2^ *−* 4**C** has a square root. The solution of the quadratic matrix equation is then:

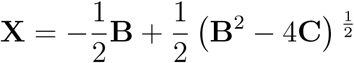

In our case, an explicit solution for **X** exists. Indeed, **X**^2^ is preceded by an identity matrix, **B** is a diagonal matrix and then always commute with **C**. Finally, as **B**^2^ and **C** are both positive semi-definite then (**B**^2^ *−* 4**C**)^1/2^ will always have a solution. Recall that

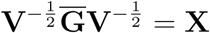

then

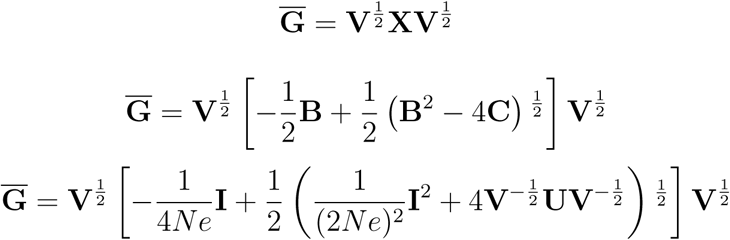

Finally by using notations (1) and (4), we obtain equation (8)

### A.2 Constraints on the orientation and shape of the G matrix in the special cases where V and M matrices have the same eigenvectors

#### A.2.1 Case without drift

Starting from equation (7)

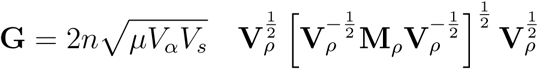

and assuming that both matrices **V**_*ρ*_ and **M**_*ρ*_ have the same eigenvectors **Q** but different eigenvalues, their spectral decompositions give **V**_*ρ*_ = **QΛ**_*s*_**Q**^*−*1^ and **M**_*ρ*_ = **QΛ**_*m*_**Q**^*−*1^ respectively, where **Λ**_*w*_ and **Λ**_*m*_ are diagonal matrices of eigenvalues. Then, equation (7) can be simplified to

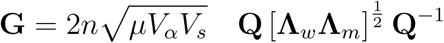

This shows that **G** matrix has the same eigenvectors **Q** as the **V** and **M** matrices, and that its eigenvalues are the geometric means of eigenvalues of **V** and **M**.

#### A.2.2 Case with drift

Starting from equation (8)

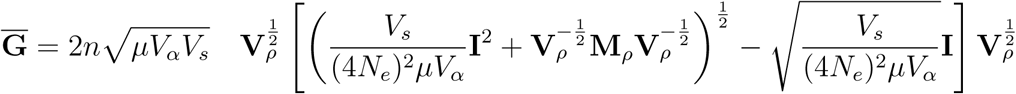

and performing a spectral decomposition of **V**_*ρ*_ and **M**_*ρ*_ as in the case without drift (see above), after simplification we o btain:

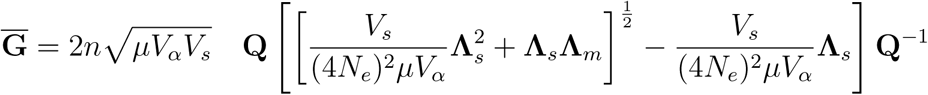

While the eigenvalues of **G** are given by the equation located between the highest level of brackets, the eigenvectors of **G** still equal **Q** the same as **V** and **M** matrices.

#### A.2.3 Graphical representation of G matrix in the special case where V and M matrices have the same eigenvectors

**Figure S1:**
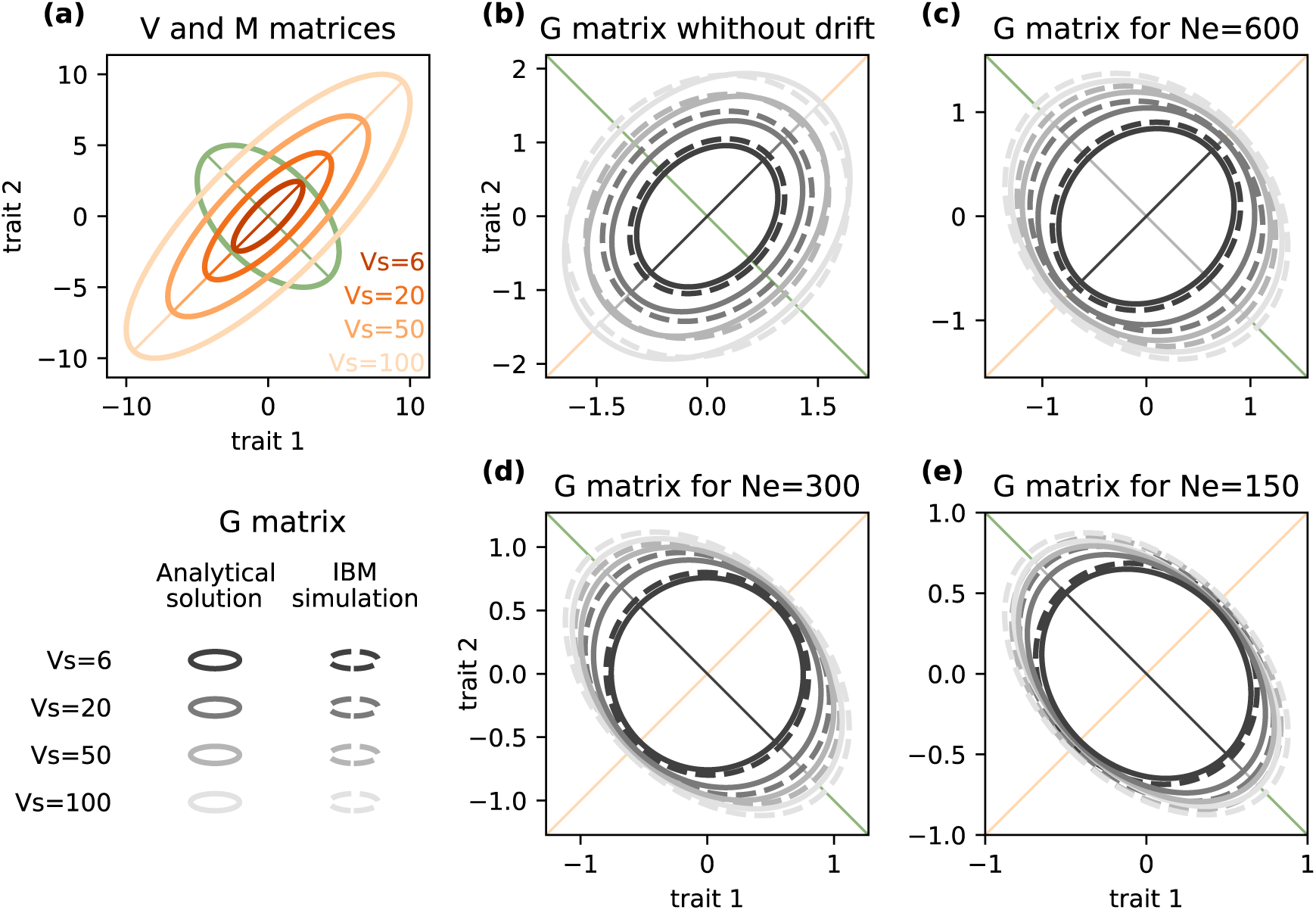
Influence of selection strength and genetic drift on the G matrix when V and M matrices have the same eigenvectors. (a) Orientation and shape of **M** and **V** matrices for respectively *ρ*_*m*_ = *−*0.5, *ϕ*_*m*_ = 1, *V*_*α*_ = 0.05, *ρ*_*s*_ = 0.7,*ϕ*_*s*_ = 1, *V*_*α*_ = 6, 20, 50, 100. (b-f) see Figure 1 for a detailed legend.

### A.3 G matrix in the House of cards regime

**Figure S2:**
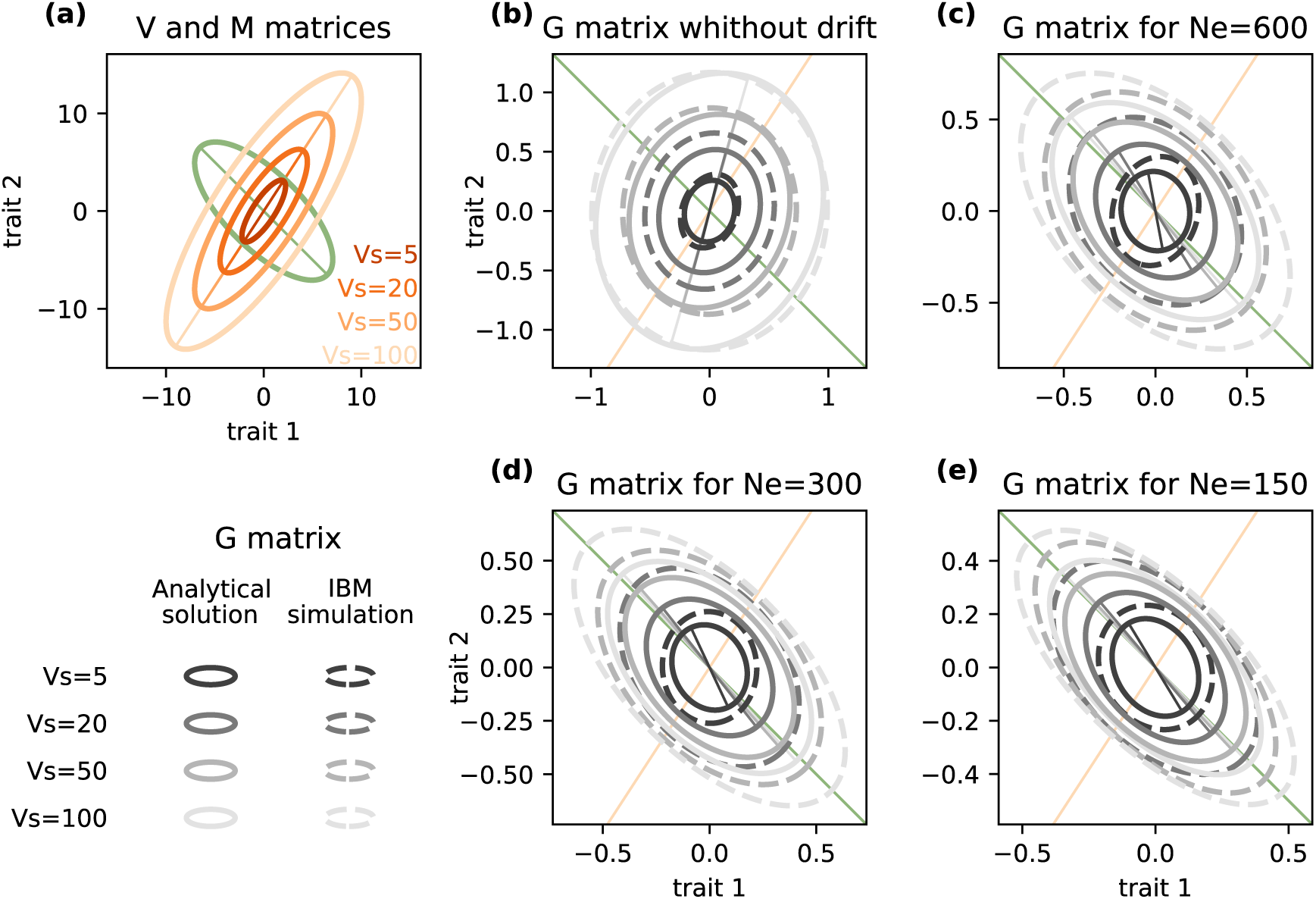
Influence of selection strength and genetic drift on the G matrix (House-of-Cards regime). (a) Orientation and shape of **M** and **V** matrices for respectively *ρ*_*m*_ = *−*0.7, *ϕ*_*m*_ = 1, *V*_*α*_ = 0.05 and *ρ*_*s*_ = 0.8, *ϕ*_*s*_ = 2, *V*_*s*_ = 5, 10, 50, 100. First eigenvectors are represented colored lines. We set n=50 and the *µ* = 0.0002. (b-f) see Figure 1 for a detailed legend.

